# Task-space dimensions guide human exploration in complex environments

**DOI:** 10.64898/2026.04.29.720265

**Authors:** Jiahui An, Jiewen Hu, Yilin Elaine Wu, Chun Liu, Siyu Ning, Fanyu Zhu, Yuhang Pan, Ruosi Wang, Ni Ji

## Abstract

Efficient exploration is essential for rapid learning in high-dimensional environments, where rewards depend on a sparse and unknown subset of task dimensions. Here, we developed a multidimensional value-learning task in which participants explored without explicit knowledge of the underlying structure. Across multiple task designs, participants consistently demonstrated successful learning under structural uncertainty. Behavioral analyses revealed that over half of participants adopted a structured exploration strategy—Full Dimension Scan (FDS)—with three key properties. First, participants systematically isolated and sampled one candidate dimension at a time, effectively reducing search complexity. Second, FDS was temporally persistent within dimensions while maintaining diverse sampling of feature values, enabling efficient local inference. Third, participants sequentially deployed FDS across dimensions, achieving broad coverage of the task space. Importantly, this strategy transferred across tasks: prior experience in lower-dimensional environments promoted stronger FDS use and improved performance in higher-dimensional settings. Consistent with these findings, a dimension-guided exploration model incorporating dimension-level attentional control best captured human behavior. Together, these results identify dimension-guided exploration as a core principle that transforms high-dimensional search into tractable, structured learning, providing new insight into adaptive behavior in complex environments.

## Introduction

Exploration is a fundamental process through which humans and other animals acquire information about unfamiliar environments and use that information to guide future behavior. This process is especially important when feedback is sparse, outcomes are uncertain, and learners must generalize from limited experience. In complex high-dimensional environments, exploration requires more than sampling unfamiliar options: learners must identify which aspects of the environment are informative, organize observations into useful representations, and infer which features or dimensions are relevant for reward. Efficient exploration may therefore depend on actively constructing informative experience and uncovering the latent structure of the task (Gershman & Niv, 2010; Collins & Frank, 2013). Understanding how humans organize this process is essential for explaining flexible learning in complex environments.

Human exploration has been extensively studied using multi-armed bandit and contextual bandit paradigms. These tasks provide tractable frameworks for examining the exploration–exploitation trade-off and have shown that human exploration can be partly explained by directed exploration, in which choices are biased toward uncertain or informative options, and random exploration, in which choice stochasticity increases under uncertainty (Daw et al., 2006; Wilson et al., 2014; Gershman, 2018; Speekenbrink & Konstantinidis, 2015). Contextual bandit and function-learning studies further show that people can generalize across contextual variables when learning cue–reward mappings (Krause & Ong, 2011; Lucas et al., 2015; Schulz et al., 2018; Wu et al., 2018; Parpart et al., 2017). However, these paradigms typically involve low-dimensional task spaces and restricted action sets, leaving open how humans organize exploration in larger environments structured by multiple latent or compositional dimensions.

Related work on multidimensional learning has shown that humans can use selective attention, rule learning, hypothesis testing, and feature-based reinforcement learning to identify reward-relevant attributes and update behavior accordingly (Kruschke, 1992; Love et al., 2004; Goodman et al., 2008; Gershman et al., 2010; Wilson & Niv, 2012; Niv et al., 2015; Mack et al., 2016; Wunderlich et al., 2011; Akaishi et al., 2016; Ballard et al., 2018; Farashahi et al., 2017; Wang & Rehder, 2017). Active learning studies further suggest that learners can select informative evidence to distinguish among candidate hypotheses (Oaksford & Chater, 1994; Nelson, 2005; Markant & Gureckis, 2014). Yet these studies have focused primarily on value updating, rule inference, or hypothesis selection, rather than on the behavioral structure of exploration itself. In many computational accounts, exploration is treated as stochastic choice noise or uncertainty-driven sampling, rather than as an organized strategy for generating informative experience.

Here, we address a fundamental question: how do humans achieve efficient exploration in high-dimensional environments when the structure of the task is unknown? We introduce a novel multidimensional value-learning paradigm that captures exploration in a large, compositional action space, allowing participants to actively construct informative experience rather than select from predefined options. This approach reveals that human exploration is not well described by randomness or uncertainty-driven sampling alone, but is instead organized around latent task dimensions. We identify a robust and previously uncharacterized strategy—Full Dimension Scan (FDS)—through which individuals systematically probe one dimension at a time, reducing the complexity of high-dimensional search. This strategy is temporally persistent, spans multiple dimensions, and transfers across tasks to support improved performance in more complex environments. A dimension-guided exploration model formalizes these findings, showing how dimension-level control can generate human-like behavior. Together, these results uncover a fundamental principle of human exploration, demonstrating how structured, dimension-wise sampling enables scalable and adaptive learning in complex environments.

## Results

### Humans learn effectively without knowledge of task dimensional structure

To study how humans explore and learn in multidimensional environments with unknown task structure, we developed a multidimensional arrangement task that expanded both the dimensionality and the action space of standard value-learning paradigms. The task retained experimental control while allowing participants to actively sample the uninstructed structure of the environment. Rather than choosing one option from a small set, participants arranged nine compound food stimuli into three rows and received score feedback after submitting each arrangement (Fig. 1a). Row position determined how strongly each item contributed to the final score: coin icons indicated the relative row weights, with items placed in upper rows contributing more than items placed in lower rows. Thus, each round required participants to decide not only which stimuli were likely to be valuable, but also how to allocate them across positions with different reward consequences.

**Fig. 1.**
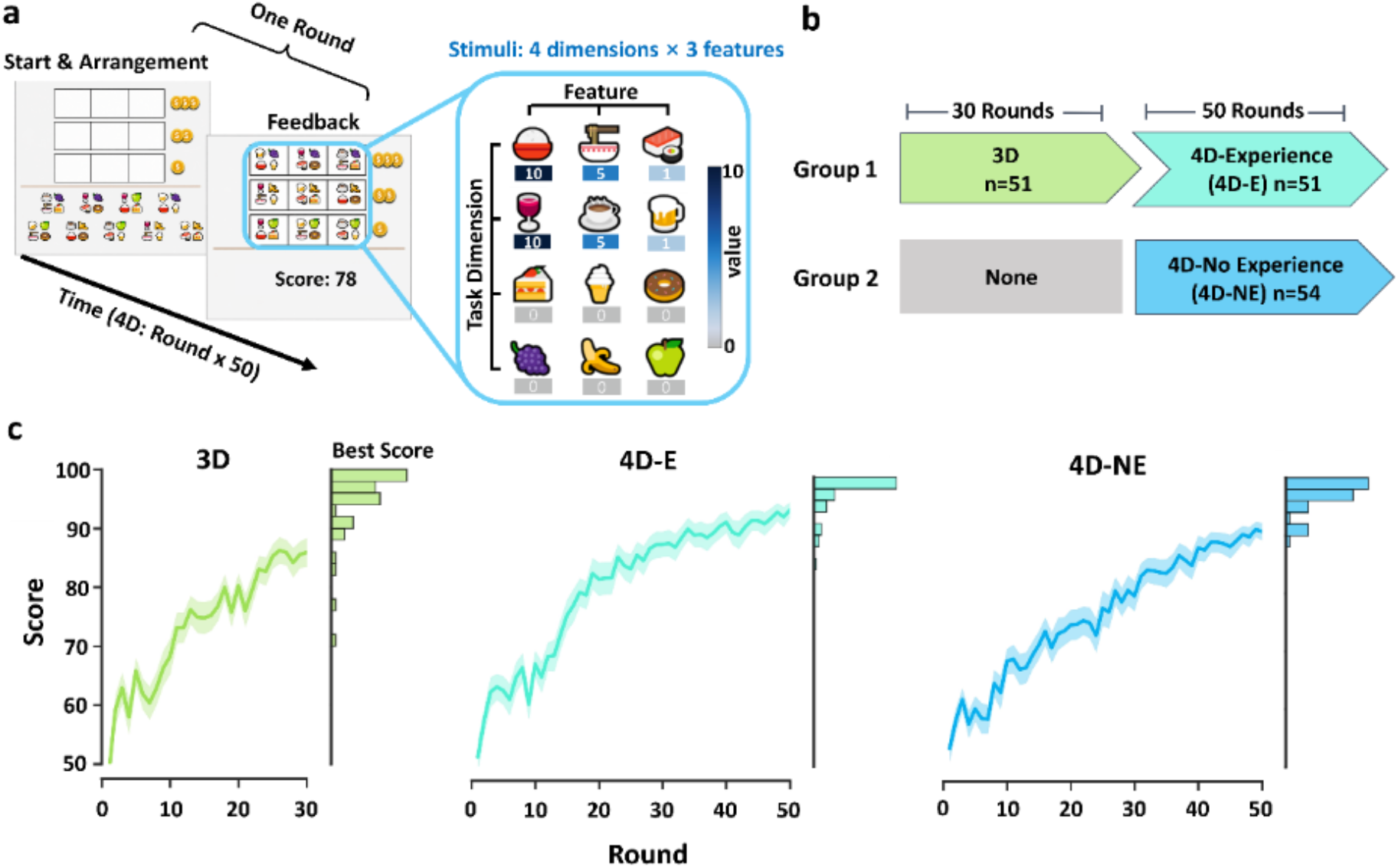
Multidimensional value-learning task and behavioral evidence of successful learning. **a**, Example round from the 4D arrangement task. Participants arranged nine compound food stimuli into three rows and received score feedback after each arrangement. Coin icons indicate the relative row weights, with upper rows contributing more to the final score. Each stimulus was defined by features from multiple task dimensions, only a subset of which was reward-relevant. **b**, Experimental design. One group completed a 30-round 3D task before entering the 50-round 4D task, whereas a separate group completed the 4D task without prior 3D experience. **c**, Task performance across groups. Line plots show mean score across rounds, with shaded bands indicating s.e.m. Marginal histograms show the distribution of each participant’s best score. Scores increased across rounds in all groups, indicating that participants learned to improve their arrangements from feedback.

We designed two task settings, a 3D task and a 4D task, which differed in the number of feature dimensions defining each stimulus. In the 3D task, each stimulus was composed of one feature from each of three dimensions; in the 4D task, each stimulus contained one feature from each of four dimensions, with three possible feature values per dimension (Fig. 1a). Critically, participants were not explicitly informed about the dimensional structure of the stimulus space, the number of reward-relevant dimensions, or which feature values within those dimensions were associated with higher reward. In addition, the reward-relevant dimensions were randomly assigned across participants. Successful performance therefore required participants to infer, from score feedback alone, both the dimensional organization of the stimulus space and the reward relevance of candidate dimensions.

To examine whether experience in a lower-dimensional task and the exploratory strategies formed in that context could transfer to a higher-dimensional task and facilitate performance in more complex environments, we employed a between-subject design. One group of participants first completed a 30-round 3D task and then proceeded to a 50-round 4D task, whereas another group directly entered the 4D task without any prior experience. Participants were randomly assigned to these 2 groups with different task procedure. This design yielded data from three conditions: the 3D condition, the 4D-Experience (4D-E) condition, and the 4D-No Experience (4D-NE) condition.

We first validated that participants could learn under this unknown multidimensional structure. Participants improved their performance across rounds in all task groups (Fig. 1c).

Mean scores increased over time in all tasks. The distribution of best scores further showed that many participants reached high-scoring arrangements during the task (Fig. 1c). Together, these learning curves and best-score distributions indicate that participants could use score feedback to improve their choices, even when both the task dimensions and the number of reward-relevant dimensions were not explicitly instructed. Having established that the paradigm elicited learning in an unknown multidimensional setting, we next asked what behavioral strategies enabled participants to search efficiently in the large arrangement space.

### A stereotyped Full Dimension Scan strategy dominates exploratory behavior

We constructed a task state space based on the structural similarity among all possible choices in the task, such that participants’ choice on a given round was represented as a choice vector embedded within this space. Based on each participant’s past choices and associated score feedback, we employed a feature-based reinforcement learning approach to estimate the optimal choice on each round, thereby obtaining a corresponding optimal choice vector within the state space.

Within this state space, we computed two metrics to quantify choice behavior. The first metric was the deviation, operationalized as the distance, between the participant’s observed choice vector and the current optimal choice vector. The second metric was the deviation, operationalized as the distance, between the participant’s observed choice vector on the current round and the participant’s observed choice vector from the immediately preceding round. We defined the former metric, the deviation from the current optimal choice vector, as the Exploration Index (EI) (Fig. 2a). Previous studies of human exploration have shown that exploratory behavior typically decreases as learning progresses. Consistent with this literature, in the 3D, 4D-E, and 4D-NE tasks, the deviation from last choice and the Exploration

**Fig. 2.**
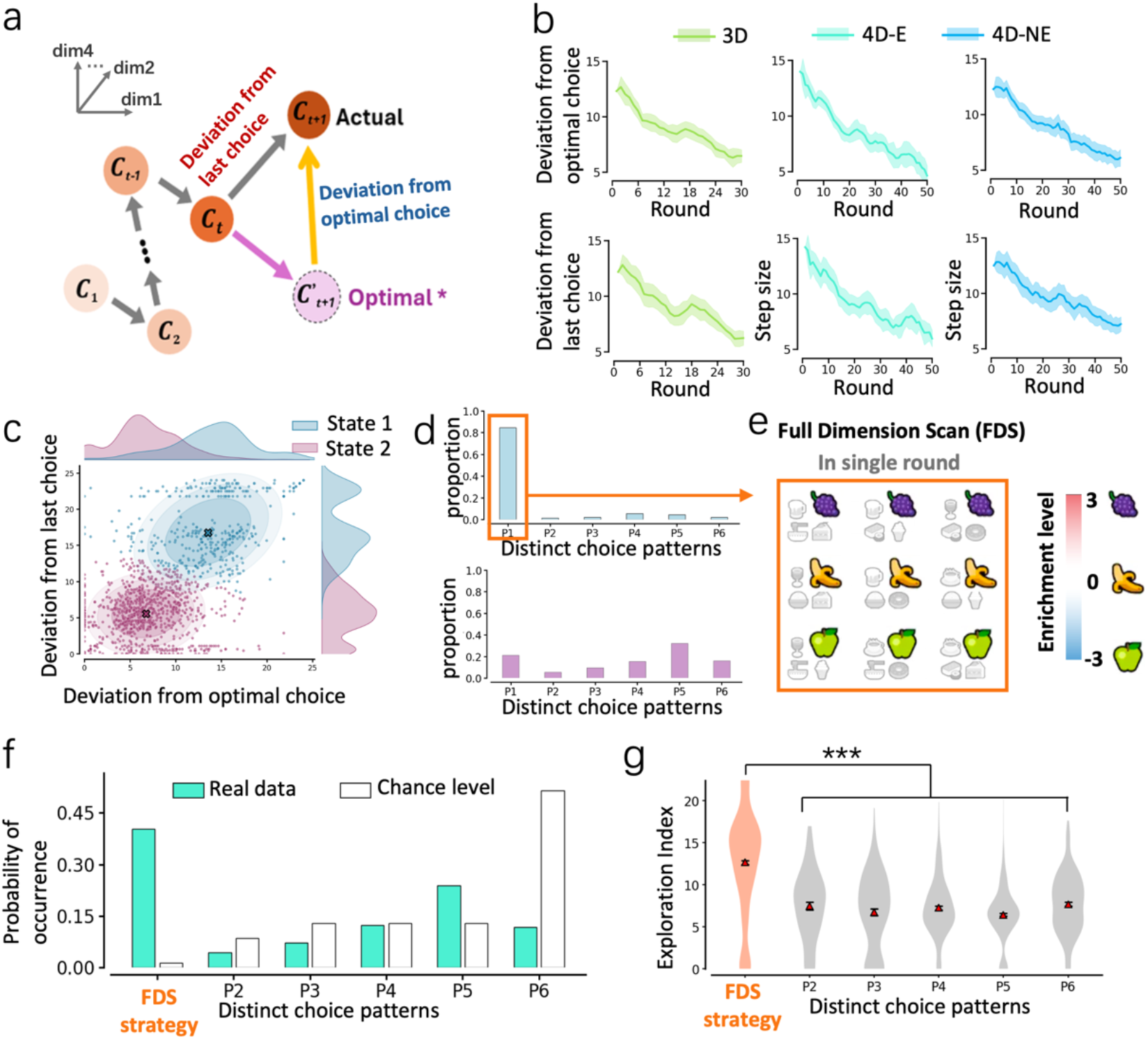
Quantification of exploration behavior and emergence of a stereotyped exploration strategy. **a**, Schematic illustration of the task state space and the two computational metrics used to quantify exploration. **b**, Temporal dynamics of exploration metrics across learning in 3D, 4D-E, and 4D-NE tasks. **c**, Behavioral state classification based on the joint distribution of the two metrics. Choices with large deviation from the last choice and high deviation from the optimal choice were classified as exploratory (State 1, purple), whereas choices with small deviation from the last choice and low deviation from the optimal choice were classified as exploitative (State 2, blue). Marginal distributions are shown on the axes. Classification was performed using Gaussian Mixture Models (GMM). **d**, Distribution of choice patterns across exploratory and exploitative states. Exploratory states were dominated by a specific structured choice pattern, whereas exploitative states showed no clear preference for any particular pattern. **e**, Schematic illustration of the Full Dimension Scan (FDS) strategy. **f**, Comparison between the observed probability of different choice patterns in human behavior and their computed chance level. **g**, Exploration Index across different choice patterns. Mean are reported as red triangles for each pattern. Two-sample Welch’s t-tests with Bonferroni correction were used to compare the FDS choice pattern against all other patterns individually. The FDS choice pattern exhibited a significantly higher Exploration Index than all other patterns. Error bars and shaded bands denote s.e.m. \**p* < 0.05, \*\**p* < 0.01, \*\*\**p* < 0.001

Index (EI) both exhibited a progressive decline over the course of the task (Fig. 2b). Using these two metrics, we classified participants’ choices on each round into two distinct states. One state was characterized by a higher EI and larger deviation from the last choice, whereas the other exhibited a lower EI and smaller deviation from the last choice. We interpreted the former as an exploratory state and the latter as an exploitative state (Fig. 2c)

We next examined the distribution of choice patterns within these states. Exploratory choices were dominated by a stereotyped structured pattern, whereas exploitative choices showed no clear preference for any choice pattern (Fig. 2d). The characteristic exploratory pattern is illustrated in Fig. 2e: within a single round, participants arranged the three features of one candidate dimension into three separate rows, repeating each feature within its corresponding row. We refer to this behavior as Full Dimension Scan (FDS).

To evaluate whether FDS reflected a genuine property of human behavior rather than a by-product of task constraints, we compared the empirical probability of each choice pattern with its computed chance level in all possible choices. FDS occurred significantly more often than expected by chance (Fig. 2f). Moreover, the Exploration Index associated with FDS was significantly higher than that of all other choice patterns (Fig. 2g). These findings identify FDS as a stereotyped exploratory strategy.

### FDS behavior is persistent within and across dimensions, while exhibiting choice diversity

We next characterized the temporal structure of FDS across multiple rounds. As illustrated in Fig. 3a, when participants adopted FDS, their exploration often persisted within the same candidate dimension while sampling diverse feature arrangements from that dimension. Participants also applied FDS across successive rounds to examine different task dimensions sequentially.

**Fig. 3.**
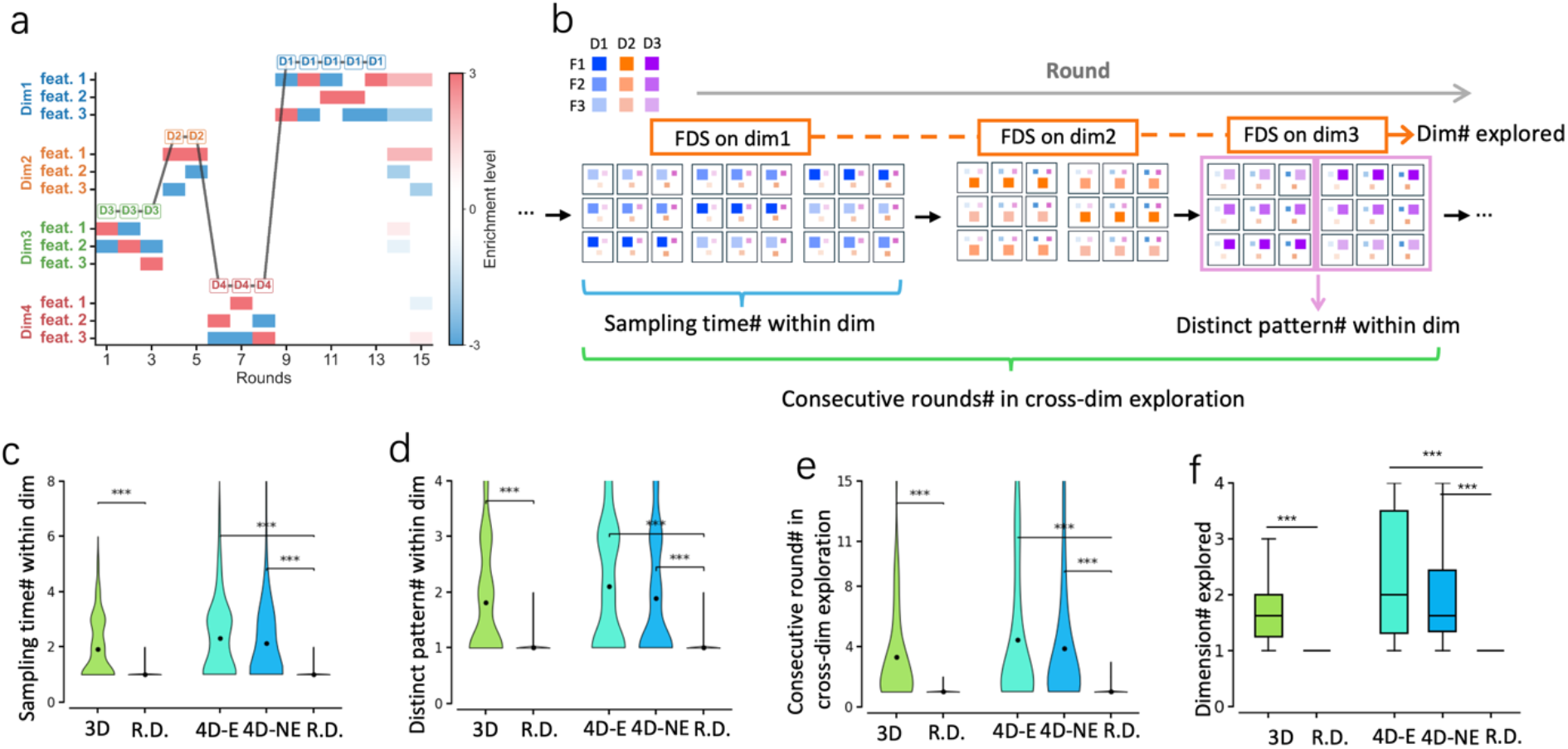
FDS behavior is persistent within and across dimensions, while exhibiting choice diversity. **a**, Example exploration trajectories illustrating the behavioral structure of FDS across rounds within a single game. **b**, Schematic illustration of the four quantitative metrics used to characterize FDS exploration at the population level. **c**, The distribution of within-dimension bout length for different task conditions and random control. **d**, Within-dimension pattern diversity for different task conditions and random control. **e**, Cross-dimension bout length for different task conditions and random control. **f**, Number of dimensions explored. Violin plots display data distribution with mean in black points. Box plots showing median, 25%-75% percentile, whiskers:1.5IQR. Real behavior (3D, 4D-E, 4D-NE) was compared to random control using two-sample t-tests (two-tailed). \**p* < 0.05, \*\**p* < 0.01, \*\*\**p* < 0.001.

To quantify these properties at the population level, we computed four metrics (Fig. 3b): (i) persistence of sampling within a single dimension, measured as the number of sampling rounds; (ii) diversity of sampling patterns within a single dimension; (iii) the number of consecutive rounds in which FDS was sustained across dimensions; and (iv) the number of distinct dimensions explored during a game.

Participants sampled within a single dimension for significantly more rounds than expected by chance (Fig. 3c). Within a sampled dimension, they also exhibited significantly greater diversity in choice patterns than expected by chance (Fig. 3d). Across rounds, participants maintained FDS for longer consecutive sequences than chance levels (Fig. 3e) and explored multiple task dimensions during a game, often approaching the total number of available dimensions and significantly exceeding chance levels (Fig. 3f).

Together, these findings revealed that participants’ exploration behavior under uninformed task environment was organized around the task dimension, exhibiting persistence both within and across task dimensions. Exploration within a given dimension showed substantial choice diversity, and participants systematically sampled across multiple task dimensions. These characteristics of FDS exploratory behavior suggest that participants did not sample randomly; rather, they leveraged the dimensional structure of the task to guide exploration in a structured and systematic manner.

### Greater use of the FDS strategy is associated with better task performance

We next examined the relationship between FDS and task performance. Across all three task phases, a higher proportion of FDS rounds was associated with higher best scores and fewer rounds required to complete the task (Fig. 4a). Participants who achieved the full score also exhibited higher FDS proportions than participants who did not (Fig. 4b). In addition, the full-score rate increased with the number of task dimensions explored using FDS (Fig. 4c). These findings indicate that greater use of FDS, particularly when deployed across multiple dimensions, was associated with more successful task performance.

**Fig. 4.**
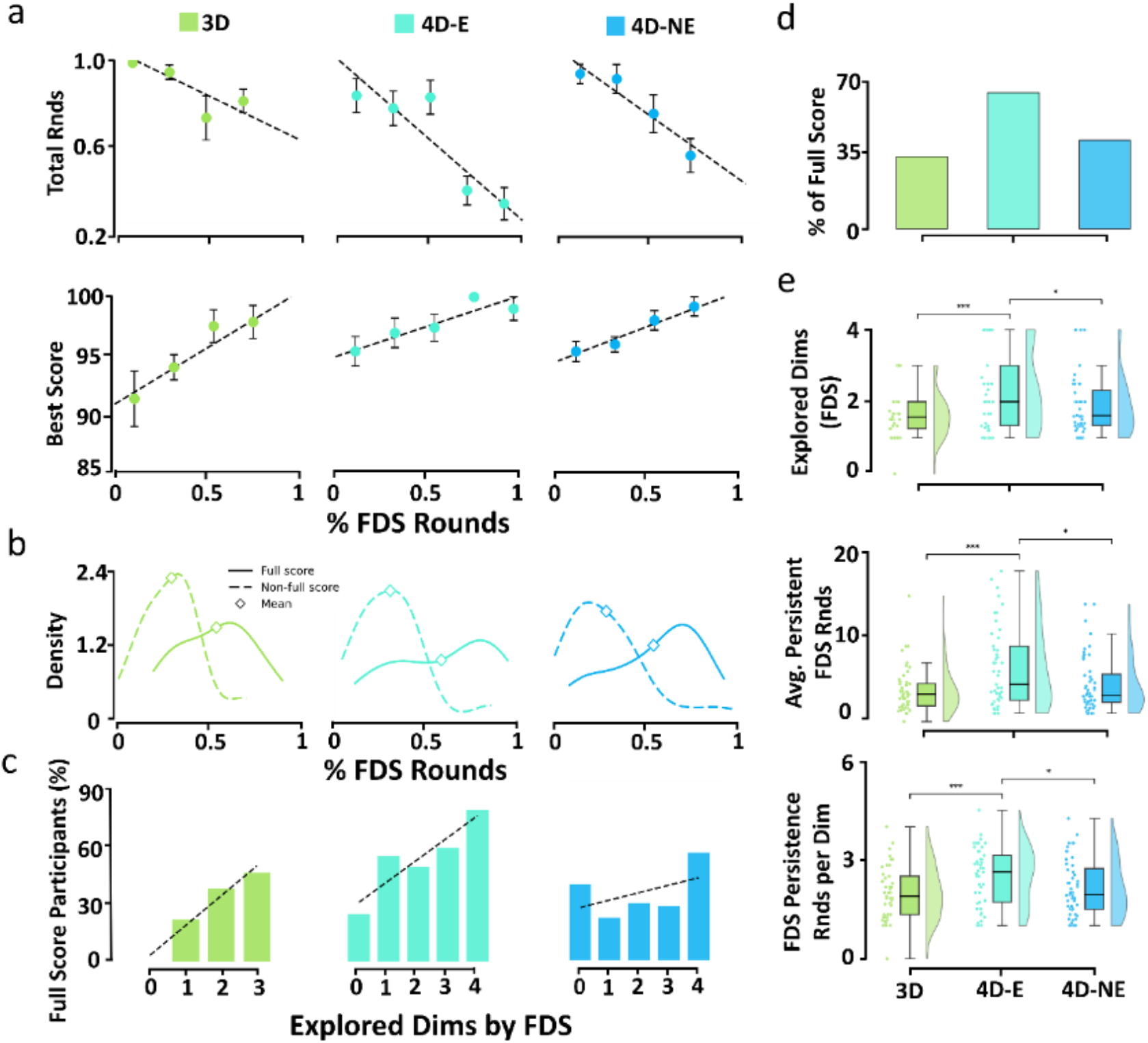
Greater use of FDS is associated with better task performance and is more strongly expressed in the experienced 4D condition. **a**, Across the 3D, 4D-E and 4D-NE tasks, the proportion of rounds in which participants used FDS was plotted against two performance measures: the number of rounds required to reach full performance (top; fewer rounds indicate better performance) and the best score achieved (bottom). In all three task conditions, greater FDS use was associated with faster attainment of full performance and higher best scores (3D: r = 0.3213, p = 0.02436; 4D-E: r = 0.4127, p = 0.002617; 4D-NE: r = 0.4631, p = 0.0007082). **b**, Distributions of FDS use in participants who did or did not achieve full score, showing that full-score participants exhibited a higher proportion of FDS use than non-full-score participants. **c**, Percentage of participants achieving full score as a function of the number of task dimensions explored using FDS, shown separately for the 3D, 4D-E, and 4D-NE tasks. In all task conditions, the proportion of full-score participants increased with the number of dimensions explored using FDS. **d**, Percentage of participants achieving full score in the three task conditions, showing a higher full-score rate in 4D-E than in 4D-NE. **e**, Comparison between 4D-E and 4D-NE participants on three FDS-related measures: the number of consecutive rounds maintaining FDS, the number of dimensions explored using FDS, and the length of FDS runs within each dimension. Participants in 4D-E showed higher values on all three measures. Box plots show the median (center line), interquartile range (25th–75th percentiles; box), whiskers extending to 1.5 times the interquartile range, and individual participants (dots); black diamonds indicate the mean. Error bars denote s.e.m. \**p*<0.05, \*\**p*<0.01, \*\*\**p*<0.001.

We then asked whether prior experience modulated performance in the 4D task. Participants showed a higher full-score rate in the 4D-Experience condition (4D-E) than in the 4D-No Experience condition (4D-NE; Fig. 4d), raising the possibility that experience from the 3D task generalized to the later 4D task. Consistent with this account, participants in 4D-E expressed stronger FDS behavior than those in 4D-NE, including longer consecutive FDS runs, exploration of more dimensions using FDS, and longer FDS runs within each explored dimension (Fig. 4e). Thus, the superior performance in 4D-E may reflect transfer of an effective exploration policy, expressed behaviorally as broader and more persistent use of FDS.

### A dimension-guided exploration (DGE) model captures human exploration better than alternative models

The behavioral analyses identified FDS as a robust strategy associated with better performance, but they did not specify the computational process that could generate such structured exploration. We therefore asked whether a model with dimension-level attentional control could account for participants’ trial-by-trial choices and reproduce the FDS patterns observed in human behavior.

To this end, we developed a dimension-guided exploration (DGE) model that combines feature-value learning with dimension-level control over exploration and exploitation (Fig. 5a). The model was designed to capture the behavioral structure suggested by the preceding analyses: FDS behavior is generated through a single-dimension attention, whereas other exploratory or exploitative choices can be guided by multi-dimension integration. At the start of each round, the model determines whether choice is guided by a single focused dimension or by integrated information across dimensions, and then generates an arrangement accordingly.

**Fig. 5.**
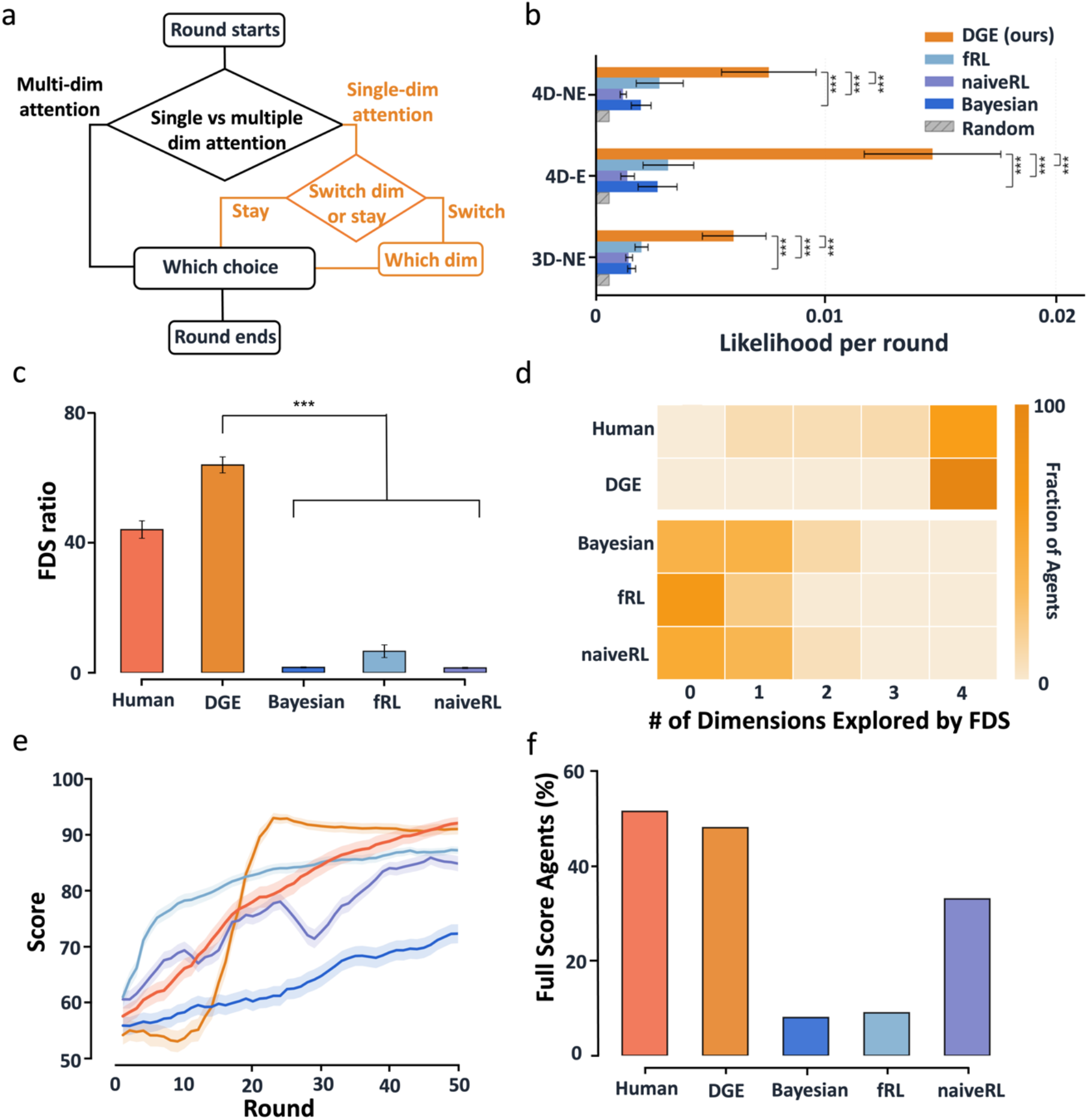
The DGE model captures human-like exploration and learning. **a**, Schematic of the dimension-guided exploration model. The model determines whether each round is guided by a single-dimension attention or by multi-dimension integration, and generates an arrangement accordingly. **b**, Model comparison based on likelihood per round. Across all task phases, the DGE model fit human choices better than several existing models. **c**, Proportion of FDS choices in humans and model agents. The DGE model reproduced the high level of FDS behavior observed in humans, whereas alternative models produced substantially fewer FDS choices. **d**, Number of dimensions explored through FDS behavior. The DGE model most closely matched the human distribution, with both humans and the DGE model frequently exploring all four dimensions in the 4D task. **e**, Learning curves of model agents across rounds. After an initial exploratory period, the DGE model showed a rapid increase in score and reached higher performance than alternative models. **f**, Full-score rate in humans and model agents. The DGE model achieved a full-score rate close to that of human participants and higher than that of alternative models. Error bars and shaded bands denote s.e.m. \**p*<0.05, \*\**p*<0.01, \*\*\**p*<0.001.

We first compared the DGE model with previously proposed models, including naive reinforcement learning (naïve RL, Niv et al., 2015) model, feature reinforcement learning (fRL, Niv et al., 2015) model, and Bayesian learning model (Song et al., 2022). Model fits were evaluated using likelihood per round. Across all task phases, the DGE model provided a better fit to participants’ choices than the alternative models (Fig. 5b). This result indicates that human behavior in the task was not fully captured by models that learned option values, feature values, or Bayesian hypotheses without explicitly modeling dimension-guided exploration.

We next examined whether the DGE model reproduced the behavioral signatures of human exploration. Human participants showed a high proportion of FDS choices, whereas the alternative models produced little FDS behavior. By contrast, the DGE model generated a level of FDS behavior comparable to that observed in humans (Fig. 5c). We further quantified the number of dimensions explored through FDS. In the 4D task, humans frequently explored all four dimensions, and the DGE model produced the most similar distribution among all models (Fig. 5d). Thus, the model captured not only the likelihood of participants’ choices, but also the characteristic pattern of dimension-wise exploration observed in human behavior.

Finally, we evaluated task performance in model agents using each model’s optimized parameter settings. The DGE model showed a distinctive learning profile: after an initial exploratory period, its score increased rapidly and reached a higher level than that of the alternative models (Fig. 5e). At the level of task success, the DGE model also achieved a full-score rate close to that of human participants and higher than that of alternative models (Fig. 5f). Together, these results suggest that dimension-level attentional control provides a compact computational account of human exploration in this task. The DGE model best captured human choices, reproduced human-like FDS behavior, and matched the distribution of explored dimensions.

## Discussion

This study examined how humans organize exploration during learning in a complex, high-dimensional environment. We found that participants did not simply sample from the large action space at random. Instead, their exploratory behavior was structured around task-space dimensions. A prominent example of this structure was Full Dimension Scan (FDS), in which participants arranged feature values from a single candidate dimension across weighted rows, allowing them to sample that dimension in a systematic way. FDS was temporally persistent, generalized across dimensions, and showed substantial within-dimension sampling diversity. Importantly, greater use of FDS was associated with better task performance, including higher best scores, faster task completion, and a higher probability of reaching the full score. These findings suggest that dimension-guided exploration is not merely a descriptive regularity in behavior, but a structured form of exploratory learning that may help people reduce complex search spaces into tractable dimension-wise sampling.

This result extends previous work on exploration in bandit and contextual bandit tasks. Standard accounts often distinguish directed exploration, in which choices are guided by uncertainty or information value, from random exploration, in which choice stochasticity increases. The present findings suggest a complementary form of structured exploration: rather than sampling isolated options, participants appeared to construct actions that selectively tested candidate dimensions. This dimension-wise strategy may be especially useful in compositional environments, where evaluating all possible actions is infeasible but the space can be decomposed into lower-dimensional structure.

The DGE model provides a computational account of this behavior by combining feature-value learning with dimension-level attentional control. Compared with alternative reinforcement-learning and Bayesian models, the DGE model better captured participants’ trial-by-trial choices and reproduced key signatures of FDS behavior. It suggests that dimension-level attention is a useful computational principle for explaining structured exploration in this task.

This dimension-guided behavior emerged despite the absence of explicit instruction about task structure, suggesting that humans can actively organize exploration around latent dimensions inferred from feedback. Our findings highlight the potential of multidimensional cognitive tasks to capture structured exploratory behavior in complex environments. By systematically varying action-space size, dimensional structure, and the statistical regularities of stimulus environments, future studies may further reveal how humans construct informative experience and adapt in high-dimensional settings. These insights could provide a foundation for developing behavioral measures of learning, cognitive flexibility, and decision-making that may ultimately be useful in translational and clinical research.

## Notes

### Competing Interest Statement

The authors have declared no competing interest.

### Summary of Updates

The language of the article has been reorganized, and more analyses have been added.

### Funding

- Supported by the Non-profit Central Research Institute Fund of Chinese Academy of Medical Sciences: 2024-RC180-02

